# The effect of short-term activation of AHL15 on long-term plant developmental change and transcriptome profile

**DOI:** 10.1101/2022.09.05.506087

**Authors:** Omid Karami, Christiaan Henkel, Remko Offringa

## Abstract

We have previously documented that overexpression of the *Arabidopsis* nuclear protein AHL15 leads to reprogramming of somatic cells to embryonic cells (Karami *et al*., 2021) and to suppression of plant ageing (Karami *et al*., 2020). Here we show that transient (4 hours) activation of overexpressed AHL15-GR in *Arabidopsis* seedlings has long-term effects on plant development. RNA sequencing analysis detected an extensive reprogramming of the transcriptome 4 hours after AHL15-GR activation, with respectively 540 and 1107 genes showing more than 2-fold up- and down-regulation. AHL15 seemed to act in a transcription level-dependent manner, activating predominantly low expressed genes and repressing mostly highly expressed genes. Rapid decondensation of heterochromatin was observed after AHL15 activation in leaf primordia and axillary meristems, indicating that the global reprogramming of the transcriptome by transient activation of this AT-Hook domain protein might be caused by extensive modulation of the chromatin configuration. We also found that co-activated or co-repressed genes were often physically linked in small chromosomal clusters, which is in line with regulation at the chromatin level.

## Introduction

The AT-HOOK MOTIF NUCLEAR LOCALIZED protein AHL15 and other members of the plant-specific AHL protein family redundantly repress the ageing pathway by delaying the juvenile to adult and the vegetative to reproductive phase transition (Rahimi *et al*., 2022). In fact, overexpression of *AHL15* was able to prolong plant longevity (Karami *et al*., 2020), resulting in the hormone-independent induction of somatic embryos on immature zygotic embryos or seedlings (Karami *et al*., 2021), and converting the monocarpic annual *Arabidopsis thaliana* into a polycarpic plant (Karami *et al*., 2020). How AHL15 can have such a dramatic effect on developmental phase transitions is one of the key questions to be addressed.

The paradigm for transcription factors is that they are nuclear proteins that bind to upstream regulatory DNA sequences and activate and/or repress transcription of target genes by respectively facilitating or inhibiting the recruitment of RNA polymerase to the transcription start site. By contrast, proteins of the plant-specific AHL family are a specific class of nuclear proteins that unlike most transcription factors, do not bind the major groove of the DNA helix, but instead interact with the narrow minor groove of DNA (Matsushita *et al*., 2007; Ng *et al*., 2009; Zhao *et al*., 2013). AHL proteins contain at least one AT-hook DNA binding motif and a Plant and Prokaryote Conserved (PPC) domain. Based on mutants and protein-protein interaction studies, the AHL family members have been proposed to bind AT-rich DNA regions as hetero-multimeric complexes that recruit other transcription factors through their interacting PPC domains (Zhao *et al*., 2013). In addition, it has been shown that AHL proteins repress transcription of several key developmental regulatory genes, possibly through modulation of the epigenetic code in the vicinity of its DNA binding regions (Lim *et al*., 2007; Ng *et al*., 2009; Yun *et al*., 2012). Some evidence has been obtained that AHL proteins function by altering the organization of the chromatin structure (Lim *et al*., 2007; Ng *et al*., 2009; Yun *et al*., 2012; Xu *et al*., 2013). Thus, different modes of transcription regulation by the AHL proteins have been described, but since this plant specific class of nuclear proteins is not well studied, their exact mode of action is still elusive.

Many AT-rich sequences in the DNA function as matrix attachment regions (MARs), which are well known to interact with the nuclear matrix, a fibrillar network of proteins inside the nucleus. Although, MARs are widely distributed in the genome, they are commonly found at the boundaries of transcription units. MARs play important roles in the higher-order organization of chromatin structure, thereby regulating gene expression (Heng *et al*., 2004; Girod *et al*., 2007; Chavali *et al*., 2011; Wilson and Coverley, 2013). The majority of the animal AT-hook motif containing DNA-binding proteins are localized to MARs where they are associated with proteins that modulate chromatin architecture (Fusco and Fedele, 2007). Also for the plant-specific AHL proteins it has been shown that they preferentially bind to the AT-rich DNA sequences of MARs (Morisawa *et al*., 2000; Fujimoto *et al*., 2004; Lim *et al*., 2007; Ng *et al*., 2009; Xu *et al*., 2013). However, a correlation between the function of AHL proteins and their localization to MARs has not been documented yet.

Here we observed that short-term activation (4 hours) of *35S::AHL15-GR* seedlings, overexpressing a fusion between AHL15 and the dexamethasone (DEX) responsive domain of the glucocorticoid receptor, resulted in long-term effects on plant development, such as delayed flowering, enhanced branching and the recurrent formation of aerial rosettes, converting monocarpic *Arabidopsis* into a polycarpic plant. In order to understand the AHL15 mode of action, we compared the chromatin configuration and transcriptome of mock- or DEX-treated *35S::AHL15-GR* seedlings by respectively nuclear staining and high throughput next generation sequencing of transcripts (Illumina RNA-Seq) (Mortazavi *et al*., 2008). Rapid decondensation of heterochromatin was observed after AHL15-GR activation in leaf primordia and axillary meristems, indicating that the observed global reprogramming of the transcriptome by transient activation of this AT-Hook domain protein might be caused by extensive modulation of the chromatin configuration.

## Results

### Short-term activation of AHL15-GR has long-term effects on plant development

We have previously shown that Arabidopsis plants that constitutively express AHL15 (35S::AHL15) or plants that express a dexamethasone (DEX) activatable version of AHL15 (35S::AHL15-GR) and were subjected to continuous DEX treatment both showed a strong delay in the juvenile-to-adult transition and flowering time (Rahimi *et al*., 2022). In contrast, *ahl* loss-of-function mutant plants showed a premature vegetative phase change and early flowering (Rahimi *et al*., 2022). Moreover, *AHL15* overexpression induced rejuvenation of axillary meristems, resulting in enhanced branching and in the production of aerial rosettes, converting monocarpic *Arabidopsis* into a polycarpic plant (Karami *et al*., 2020).

For typical transcription factors it has been shown that they act transiently, whereas other nuclear factors have a more long-lasting effect on the gene expression profile of a cell, as they act on the chromatin structure by inducing epigenetic changes (Bratzel and Turck, 2015). In order to analyses the mode of action of AHL15, 5-day-old *35S::AHL15-GR* seedlings were submerged in water with 20 μM DEX or without DEX (mock) for 4 and 8 hours, and subsequently DEX was removed by washing the seedlings in water and transferring them to soil. DEX-incubated seedlings developed much slower and the resulting plants showed a significant delay in flowering compared to the plants derived from the mock-treated seedlings (Fig. 1A). DEX-treatment for 8 hours enhanced the phenotypes observed in the 4 hour DEX-treated seedlings and derived plants (not shown). Also, 35-day-old flowering *35S::AHL15-GR* plants that were only sprayed once with 20 μM DEX developed aerial rosettes from their axillary meristems 7-10 days after spraying (Fig. 1B), and new aerial rosettes continued to develop for at least 4 months after spraying (Fig. 1C-E). These data indicates that short-term activation of AHL15 leads to long-term changes in development, which would be in line with a function for AHL15 in chromatin remodeling.

**Figure 1.**
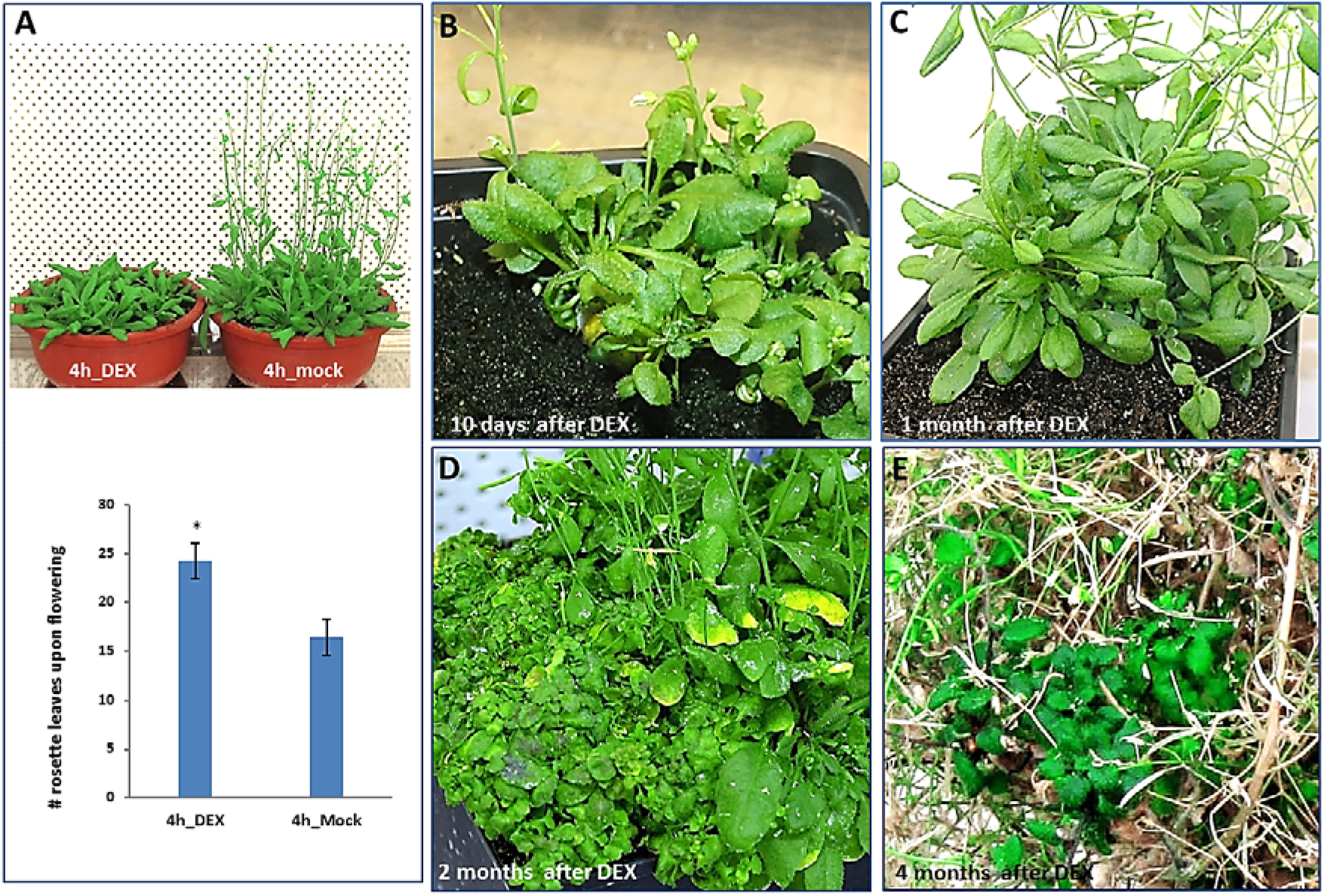
Short term activation of AHL15 has long term effects on plant development. (A) Phenotype (upper panel) and quantification of the number of rosette leaves (lower panel) of 35 day old flowering *35S::AHL15-GR* plants that were grown from 5 day old seedlings that were submerged for 4 hours in water (4h_mock) or in 20 μM DEX solution (4h_DEX). Asterisk in the graph indicates a significant difference between mock- and DEX-treated plants (Student’s *t*-test, p < 0.01), and error bars indicate standard error of the mean (n =20). (B E) Aerial rosettes developing from axillary meristems in *35S::AHL15-GR* plants 10 days (B), or one (C), two (D) or four months (E) after spraying 1 month old flowering *35S::AHL15-GR* plants with 20 μM DEX solution. Plants were grown under LD.

### AHL15 extensively reprograms the *Arabidopsis* transcriptome

In order to uncover the molecular basis of these long term developmental changes induced by AHL15, the genome-wide expression changes were compared between DEX-treated and untreated *35S::AHL15-GR* seedlings. As the major developmental changes induced by AHL15 overexpression were observed in the shoot (ref), we isolated RNA from the shoot part (i.e. shoot apex, cotyledons and top part of hypocotyl) of 5-day-old *35S::AHL15-GR* seedlings submerged for 4 hours in mock or in 20 μM DEX, or for 8 hours in 20 μM DEX. RNA sequencing was performed on RNA isolated from three biological replicates for each treatment. A slight but significant reduction of *AHL15-GR* expression was observed in DEX-treated seedlings compared to untreated samples (Table 1), suggesting that AHL15 modulates the *35S* promoter activity in this time period.

**Table 1.** Overview of raw data obtained by Illumina sequencing on RNA isolated from the shoot part of 5-day-old *35S::AHL15-GR* seedlings that were treated with water for 4 hours (4h_mock)) or with 20 μM DEX for 4 or 8 hours (resp. 4h_DEX or 8h_DEX).

| Sample # | Treatment | Raw sequencing reads | Expression level of *AHL15*<sup>1</sup> |  |
| --- | --- | --- | --- | --- |
| 1 | 4h_mock | 14285557 | 678.13 | 687.06 ± SE |
| 2 | 4h_mock | 17597921 | 636.59 |  |
| 3 | 4h_mock | 27463033 | 746.46 |  |
| 4 | 4h_DEX | 32465125 | 519.36 | 519.70 ± SE * |
| 5 | 4h_DEX | 21100996 | 526.87 |  |
| 6 | 4h_DEX | 28328909 | 512.89 |  |
| 7 | 8h_DEX | 30340913 | 519.13 | 532.90 ± SE * |
| 8 | 8h_DEX | 26594329 | 538.09 |  |
| 9 | 8h_DEX | 30496134 | 541.48 |  |

<sup>1</sup> Normalized Reads Per Kilobase Million (RPKM) values. Right column indicates the average value of the three biological replicates ± standard error of the mean.
\* Significantly different from 4h\_mock (Student's *t*-test, $p < 0.001$ )

Annotation of the reads using the *Arabidopsis* Information Resource (www.arabidopsis.org) showed that the expression of 22570 genes could be detected in the 4 hours mock samples, which is 83% of the total gene number (Table S1). Statistical analysis showed that 13483 genes were differentially expressed in 4 hours DEX-treated compared to mock treated *35S::AHL15-GR* shoot organs (Table S2). However, only 1663 genes showed a fold change of ≥ 2 (and P ≤ 0.05) after 4 hours DEX treatment (Fig. 2A, B, Table S3 and S4). Verification of the change in expression for 4 selected genes by qRT-PCR analysis showed a good agreement with the RNA-sequencing results (Table 2).

**Figure 2.**
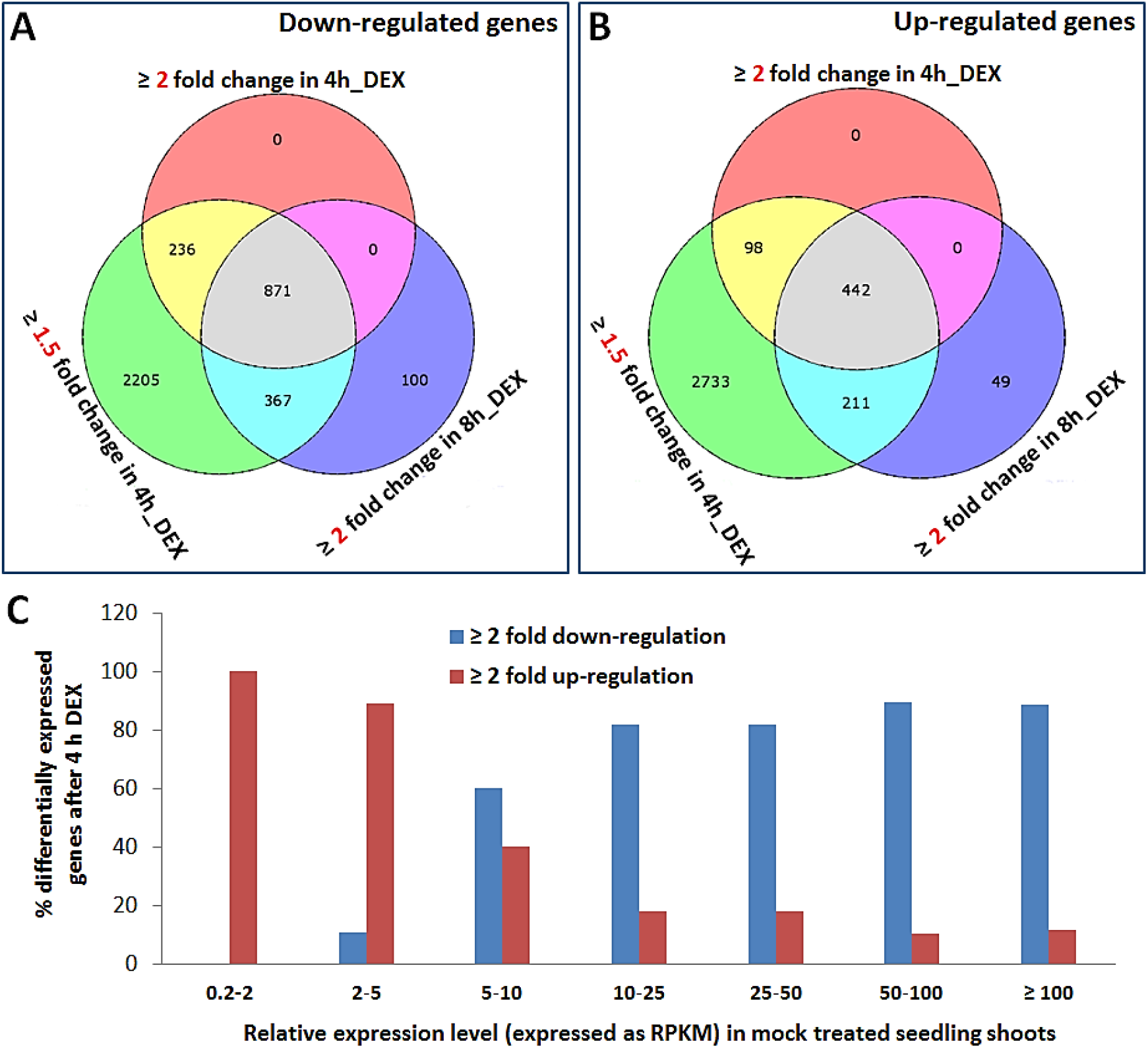
Changes in the transcriptome of *Arabidopsis 35S::AHL15-GR* seedling shoots after DEX treatment. (A, B) Venn diagrams indicating the number of (overlapping) down-regulated (A) or up-regulated (B) genes in *35S::AHL15-GR* seedling shoots with a ≥1.5 fold or ≥2 fold change in gene expression following 4 or 8 hours of DEX treatment. (D) Graph showing the percentage of genes having a specific expression level (in RPKM) in mock treated *35S::AHL15-GR* seedling shoots that are either ≥2 fold up- (red) or ≥2 fold down- (blue) regulated by 4 hours DEX treatment.

**Table 2.** Confirmation of RNA-seq data by qRT-PCR analysis.

| Locus | Name and short description | Fold change found by RNA Seq | P Value | Fold change detected by qRT-PCR<sup>1</sup> | P Value |
| --- | --- | --- | --- | --- | --- |
| AT5G59490 | Haloacid dehalogenase-like hydrolase (HAD) superfamily protein | 135.5 | 2.07E-145 | 78.30 ± 7.80 | 0.003 |
| AT4G22770 | AT hook motif DNA-binding family protein 2 | 15.19 | 8.02E-139 | 9.32 ± 1.45 | 0.005 |
| AT2G17740 | Cysteine/Histidine-rich C1 domain family protein | 0.002 | 1.21E-62 | 0.008 ± .002 | 0.0003 |
| AT1G78580 | Trehalose-6-phosphate synthase 1 | 0.35 | 2.11E-54 | 0.43 ± 0.06 | 0.021 |

<sup>1</sup> Shown is the fold changes ± standard error of the mean (n=3). P value: Student's *t*-test.

Comparison of the up- and downregulated gene sets showed a considerable overlap between 4 and 8 hours DEX treatment (Fig. 2A,B). Still, however, the number of up- (Table S5) and down-regulated (Table S6) genes increased significantly after 8 hours DEX treatment (Fig. 2A,B). Several of the genes that did not show a ≥ 2 fold change after 4 hours DEX treatment reached this level after 8 hours treatment (Fig. 2A,B). Conversely, some genes that reached ≥ 2 fold change after 4 hours, did not appear anymore in the ≥ 2 fold gene set after 8 hours (Fig. 2A,B), indicating time-dependent transcriptome changes by AHL15-GR. The latter result would be more in line with a role for AHL15 as a gene-specific transcription factor.

Notably, we found that the down-regulated genes were generally expressed at higher levels in the 4h_mock control sample, whereas the up-regulated genes were expressed at relatively low levels in the 4h_mock control sample (Fig. 2C). This data suggests that AHL15 reverses phase changes by modulating gene activity in a transcription level-dependent manner, repressing genes that are highly expressed and activating genes that are low expressed during a specific developmental phase. Global prediction of the transcriptome changes using gene ontology (GO) examination (Du *et al*., 2010) (http://bioinfo.cau.edu.cn/agriGO/) and TAIR as reference for the annotation of *Arabidopsis* genes indicated that up- and down-regulated gene sets grouped in many different biological categories (Tables S7 and S8). These results suggested that, beside its action as regulator of individual genes, AHL15 contributes to a more global reprograming of biological processes by inducing global changes in gene transcription.

Whereas most gene families represented in the transcriptome profiles (Table S2) had gene members that were either up- or down-regulated, members of the *CYSTEIN-RICH RECEPTOR-LIKE KINASE* (*CRK*) gene family only showed down-regulation of gene expression by AHL15 (Table 3). Notably, several of the down-regulated *CRKs* by AHL15 are neighboring genes that are located in a tandem arrays on chromosome 4 (Fig, 3A). This triggered us to look into the localization of up- or down-regulated genes, and surprisingly, a high rate (~75%) of co-activation or co-repression of neighboring genes by AHL15 was observed across the genome (Tables S9, and Fig. 3B). These observations suggest that AHL15 modulates gene expression in a chromosomal position-rather than a gene-specific manner.

**Table 3.** Members of the *Cysteine-rich receptor-like kinase* gene family were all down-regulated in *35S::AHL15-GR* seedling shoots after 4 hours DEX treatment

| Locus | Name and short description | Fold change | P Value |
| --- | --- | --- | --- |
| AT4G23180 | cysteine-rich RLK (RECEPTOR-like protein kinase) 10 | 0.34 | 5.49E-29 |
| AT4G23190 | cysteine-rich RLK (RECEPTOR-like protein kinase) 11 | 0.32 | 4.58E-39 |
| AT4G23200 | cysteine-rich RLK (RECEPTOR-like protein kinase) 12 | 0.04 | 1.92E-87 |
| AT4G23210 | cysteine-rich RLK (RECEPTOR-like protein kinase) 13 | 0.08 | 1.28E-44 |
| AT4G23260 | cysteine-rich RLK (RECEPTOR-like protein kinase) 18 | 0.138 | 9.12E-165 |
| AT4G23270 | cysteine-rich RLK (RECEPTOR-like protein kinase) 19 | 0.82 | 0.0098589 |
| AT1G70520 | cysteine-rich RLK (RECEPTOR-like protein kinase) 2 | 0.40 | 5.80E-33 |
| AT4G23290 | cysteine-rich RLK (RECEPTOR-like protein kinase) 21 | 0.11 | 6.19E-110 |
| AT4G23300 | cysteine-rich RLK (RECEPTOR-like protein kinase) 22 | 0.17 | 1.37E-106 |
| AT4G21400 | cysteine-rich RLK (RECEPTOR-like protein kinase) 28 | 0.77 | 7.47E-05 |
| AT4G21410 | cysteine-rich RLK (RECEPTOR-like protein kinase) 29 | 0.94 | 0.33050 |
| AT1G70530 | cysteine-rich RLK (RECEPTOR-like protein kinase) 3 | 0.40 | 7.76E-26 |
| AT4G11460 | cysteine-rich RLK (RECEPTOR-like protein kinase) 30 | 0.31 | 5.07E-19 |
| AT4G04490 | cysteine-rich RLK (RECEPTOR-like protein kinase) 36 | 0.14 | 1.29E-53 |
| AT4G04570 | cysteine-rich RLK (RECEPTOR-like protein kinase) 40 | 0.30 | 4.74E-63 |
| AT4G00970 | cysteine-rich RLK (RECEPTOR-like protein kinase) 41 | 0.09 | 2.31E-62 |
| AT5G40380 | cysteine-rich RLK (RECEPTOR-like protein kinase) 42 | 0.25 | 1.70E-23 |
| AT4G23130 | cysteine-rich RLK (RECEPTOR-like protein kinase) 5 | 0.19 | 5.63E-41 |

**Figure 3.**
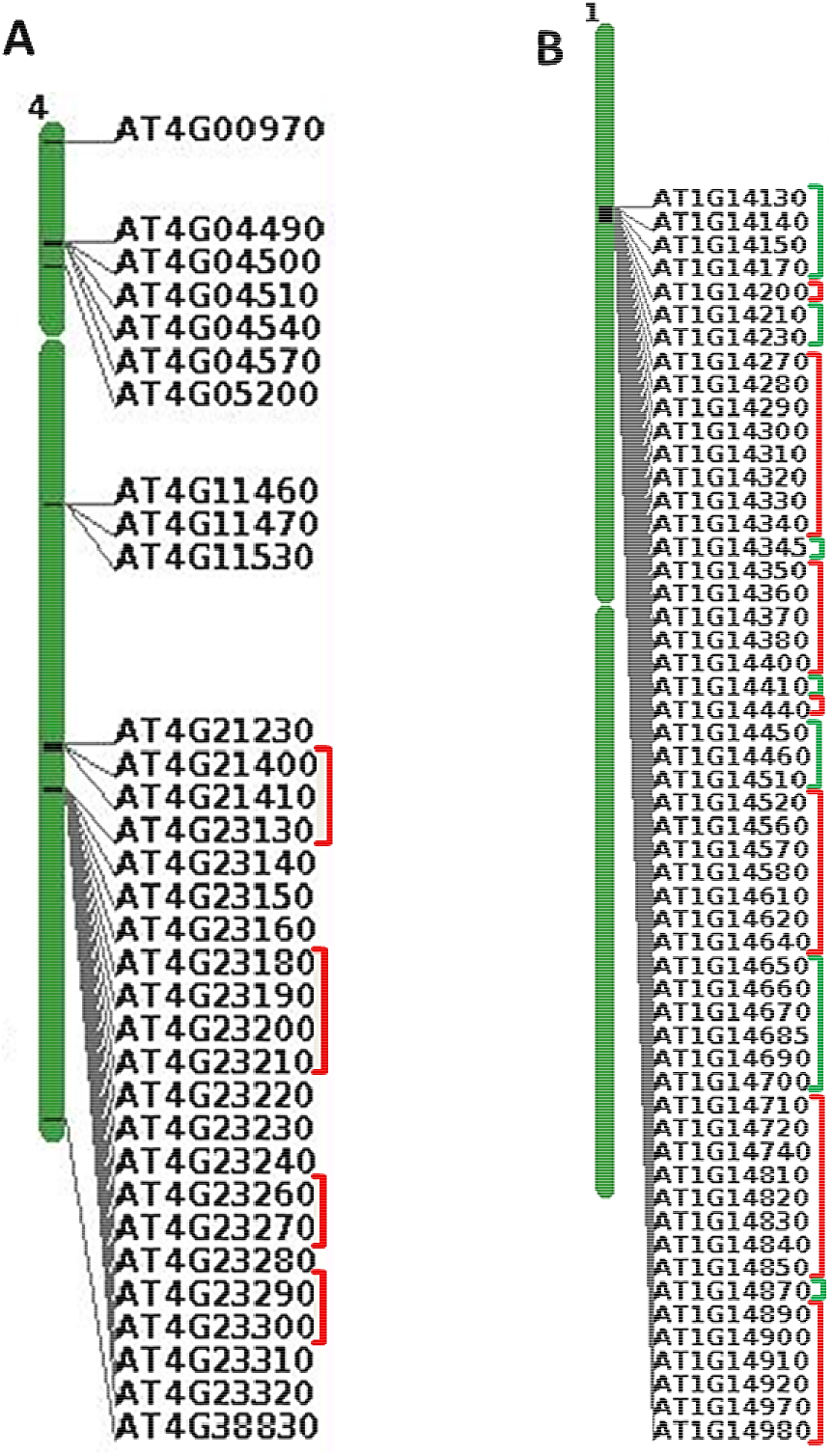
AHL15 modulates gene expression in a chromosomal region-dependent manner. (A) Schematic representation of *Arabidopsis* chromosome 4 showing the tandem arrays of *CRKs* genes on this chromosome. RNA Seq analysis on 5 day old *35S::AHL15-GR* seedlings shoots treated for 4 hours with 20 uM DEX shows that some neighboring *CRK* genes are co-repressed (indicated by the red line). (B) Schematic representation of *Arabidopsis* chromosome 1 showing for one region that neighboring genes are co-repressed (red line) or co-induced (green line) in 5 day old *35S::AHL15-GR* seedling shoots following 4 hours DEX treatment.

### *AHL15* overexpression results in reduced heterochromatin condensation

Based on previous analysis in animals that AT-hook proteins are able to change the higher order compaction of chromatin organization (Catez *et al*., 2004; Kishi *et al*., 2012; Postnikov and Bustin, 2016), we compared the chromatin organization in leaf cells of two week old *35S::AHL15* seedlings with that in wild-type leaf cells. By imaging DAPI stained nuclei or nuclei of leaf cells expressing GFP-tagged histone H2B, a considerable reduction of heterochromatin could be detected in *35S::AHL15* leaf cells compared to wild-type cells (Fig. 4A). This reduction in heterochromatin by AHL15 was even more clearly observed by imaging GFP-H2B in leaf primordia produced from axillary meristems on inflorescences (Fig. 4B). Time-lapse imaging of GFP signals revealed a rapid gradual reduction in heterochromatin condensation in DEX-treated compared to mock-treated *35S::AHL15-GR* leaf primordia (Fig. 4B), indicating that the AHL15-induced heterochromatin opening occurs in a time-dependent manner and within the time frame of the transcriptome analysis. These data support the view that the observed global reprogramming of the transcriptome by AHL15 might be caused by extensive modulation of the chromatin configuration.

**Figure 4.**
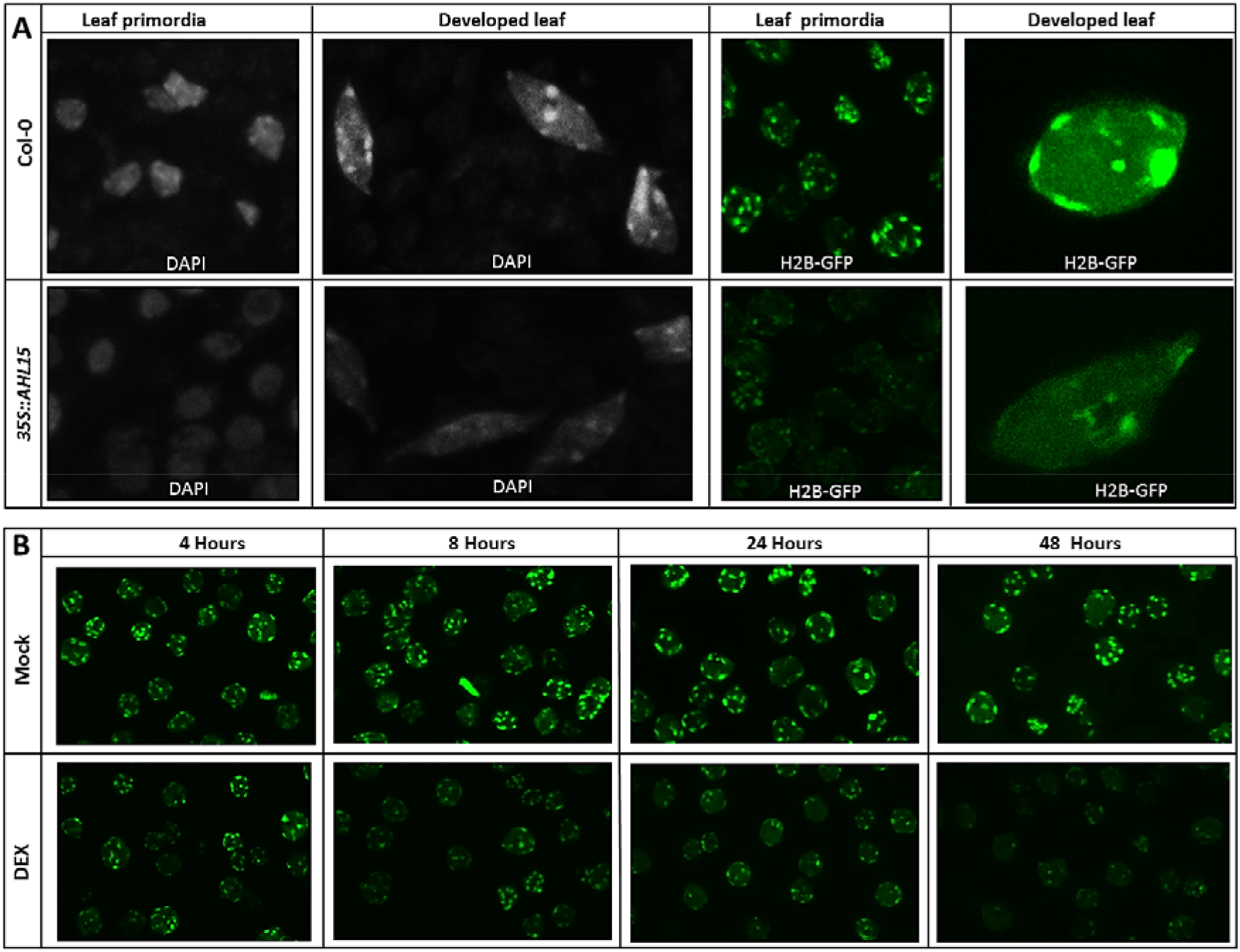
*AHL15* overexpression induces rapid heterochromatin decondensation. (A) Visualization of heterochromatin using DAPI-staining or H2B-GFP labelling of nuclei in leaf primordia or fully developed leaf cells in 2-week-old wild-type (upper row) or *35S::AHL15* (lower row) plants. (B) Heterochromatin labelling by H2B-GFP in nuclei of *35S::AHL15-GR* axillary leaf primordium cells at 4, 8, 24, or 48 hours after mock (upper row) or DEX treatment (lower row).

### Reduced photosynthesis in *35S::AHL15* plants causes sucrose-dependent seedling growth

In the transcriptome profile of DEX-treated *35S::AHL15-GR* seedlings we detected down-regulation of several photosynthesis-related genes (Table 4). The MapMan hierarchical ontology software (Thimm *et al*., 2004) showed that several of the down regulated genes encode for components of photosystem I and II, such as the light harvesting complexes, and the reaction centers (Fig. 5A, Table 4). These are among the most highly expressed genes in plants, and the fact that they are repressed by AHL15 is in line with the previously proposed transcription level-dependent manner of regulating gene expression by AHL15.

**Figure 5.**
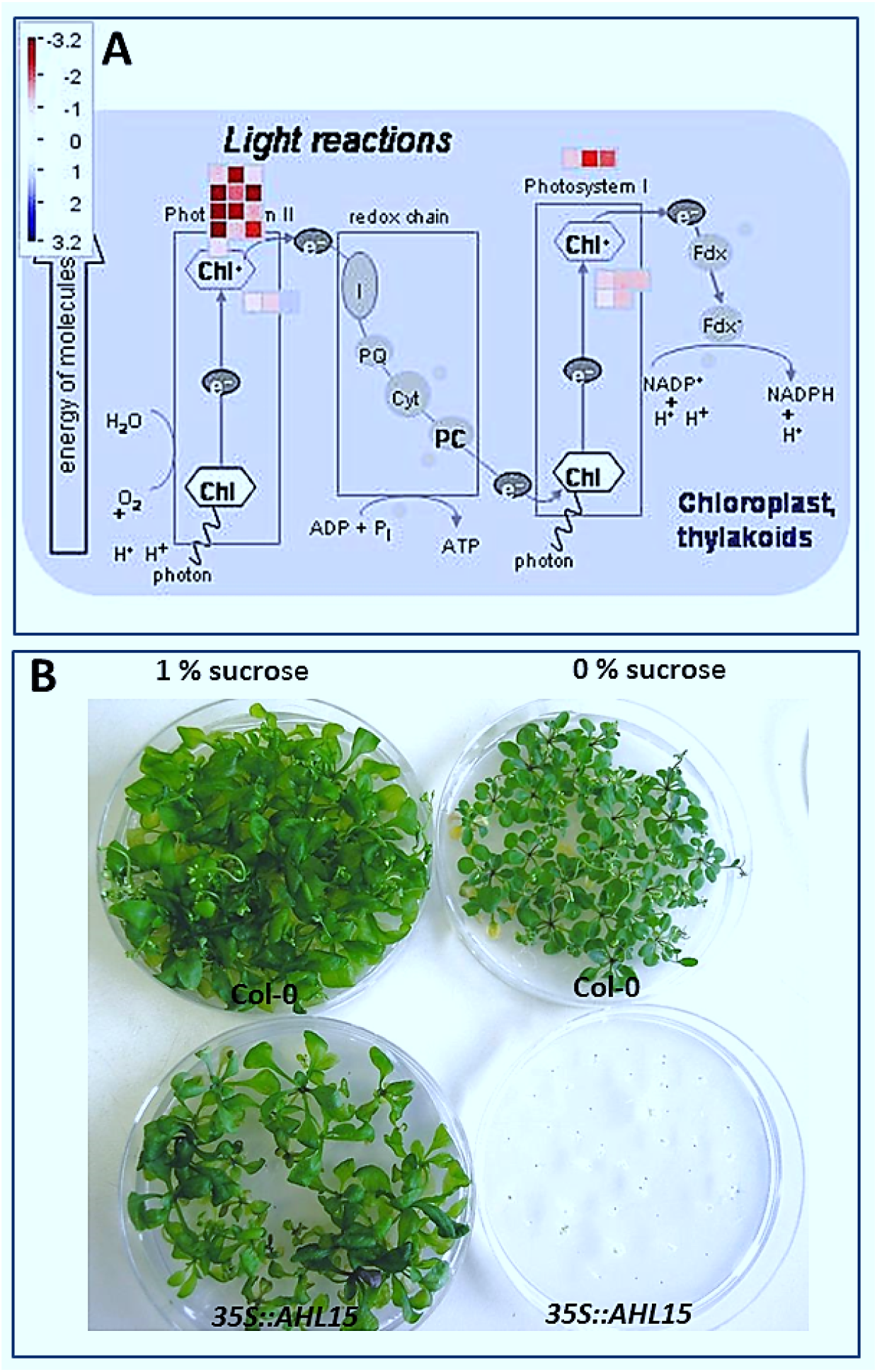
Repression of photosynthesis by AHL15 results in sucrose-dependent seedling growth. (A) A simplified scheme generated by the MAPMAN software (Thimm et al., 2004) showing that the expression of most genes encoding Photosystem I and II components is repressed in *35S::AHL15-GR* seedling shoots following 8 hours of DEX treatment. Blue squares indicate up-regulated genes, red squares indicate down-regulated genes, and gray dots indicate genes for which the expression did not change. (B) Phenotypes of 4 week old wild-type (Col-0, top) and *35S::AHL15* plants (bottom) germinated and grown on medium containing 1% sucrose (left) or no sucrose (right).

**Table 4.** List of photosynthesis-related genes that were down-regulated in *35S::AHL15-GR* seedling shoots after 8 hours DEX induction

| Locus | Name and descriptions | Expression level in 4h_water | Fold change | P Value |
| --- | --- | --- | --- | --- |
| AT4G10340 | light harvesting complex of photosystem II 5 | 3485 | 0.37 | 1.58E-44 |
| AT5G01530 | light harvesting complex photosystem II | 2405 | 0.45 | 3.93E-35 |
| AT3G08940 | light harvesting complex photosystem II | 1043 | 0.34 | 2.54E-37 |
| AT1G15820 | light harvesting complex photosystem II subunit 6 | 2323 | 0.44 | 1.29E-32 |
| AT5G54270 | light-harvesting chlorophyll B-binding protein 3 | 3520 | 0.18 | 5.28E-78 |
| AT3G47470 | light-harvesting chlorophyll-protein complex I subunit A4 | 3885 | 0.18 | 2.13E-93 |
| AT2G34430 | light-harvesting chlorophyll-protein complex II subunit B1 | 2402 | 0.08 | 1.22E-118 |
| AT3G54890 | photosystem I light harvesting complex gene 1 | 3839 | 0.23 | 2.29E-103 |
| AT1G61520 | photosystem I light harvesting complex gene 3 | 2957 | 0.39 | 8.23E-42 |
| AT2G05100 | photosystem II light harvesting complex gene 2.1 | 1773 | 0.04 | 2.11E-109 |
| AT2G05070 | photosystem II light harvesting complex gene 2.2 | 1495 | 0.05 | 7.91E-101 |
| AT3G27690 | photosystem II light harvesting complex gene 2.3 | 362 | 0.02 | 8.19E-97 |
| AT2G34420 | photosystem II light harvesting complex gene B1B2 | 5162 | 0.36 | 2.86E-51 |
| AT1G03130 | photosystem I subunit D-2 | 612 | 0.36 | 1.03E-32 |
| AT1G31330 | photosystem I subunit F | 3256 | 0.41 | 1.78E-40 |
| AT3G50820 | photosystem II subunit O-2 | 713 | 0.41 | 4.87E-38 |
| AT1G08380 | photosystem I subunit O | 3446 | 0.37 | 2.84E-58 |
| AT5G64040 | photosystem I reaction center subunit PSI-N, chloroplast | 2222 | 0.37 | 2.58E-37 |
| AT4G28750 | Photosystem I reaction center subunit IV / PsaE protein | 1711 | 0.49 | 2.94E-26 |
| AT2G30570 | photosystem II reaction center W | 2243 | 0.48 | 5.79E-26 |

Recent studies suggest that the increasing photosynthesis efficiency and sugar concentration in shoot organs of the developing young plant promote the vegetative phase change (Matsoukas *et al*., 2013; Yang *et al*., 2013; Yu *et al*., 2013). The suppression of ageing in *35S::AHL15* seedlings could therefore at least in part be caused by a delayed increase in the photosynthesis efficiency. To confirm that *AHL15* overexpression reduces the photosynthesis efficiency and thus the sugar concentration, we germinated wild-type or *35S::AHL15* seedlings on medium with or without sucrose. On medium with sucrose, both wild-type and *35S::AHL15* seedlings developed, albeit that development of *35S::AHL15* seedlings was retarded. On medium without sugar wild-type seedlings also developed, but generally much slower than sucrose grown seedlings, suggesting that photosynthesis under these *in vitro* conditions is rate limiting (Fig. 5B). In contrast, *35S::AHL15* seeds did germinate on medium without sucrose but seedling development was completely arrested (Fig. 5B). The results indicate that the germinating seedlings lack endogenous sugars, and therefore are completely dependent for their development on externally provided sucrose, most likely due to *AHL15*-mediated repression of photosynthesis during germination.

## Discussion

The *Arabidopsis* AT-hook motif nuclear-localized protein AHL15 delays phase transitions during plant development. In fact, overexpression of this protein can even reverse these phase transitions, resulting in the 2,4-D-independent induction of somatic embryos on cotyledons of immature zygotic embryos or seedlings, or in the appearance of juvenile aerial rosettes from axillary meristems of flowering plants. In addition to these initial results (Chapters 2 and 3 of this thesis), we observed that short transient activation of a constitutively expressed AHL15-GR fusion led to long-term effects on plant developmental timing and ageing. Activation of AHL15-GR by DEX treatment of seedlings resulted in a significant delay of development and flowering for up to a month after treatment, whereas spraying flowering plants with DEX resulted in the development of aerial rosettes from axillary meristems for up to 4 months after treatment. These results suggested that AHL15 establishes a longterm molecular memory, which made us wonder about the mode of action of this plant-specific class of nuclear DNA binding proteins.

Genome-wide analysis of transcriptome changes following transient activation of AHL15-GR showed that AHL15 modulates the expression level of a large number of genes participating in various biological processes. Some genes were induced after 4 hours but showed basal expression levels again after 8 hours, suggesting the direct transient activation of target genes that is typical for normal transcription factors. On the other hand, the large number of genes with a changed expression profile suggested a more global reprogramming of cellular processes. Previous studies have demonstrated that the dynamics of higher-order chromatin organization plays a critical role in both the global regulation of the transcriptome (Ho and Crabtree, 2010; Li and Reinberg, 2011; Pombo and Dillon, 2015) and the establishment of long-term molecular memory (He and Amasino, 2005; Jarillo *et al*., 2009; Harmston and Lenhard, 2013).

An additional result supporting a regional rather than single gene regulation mode was the observation that neighboring genes are co-activated or co-suppressed by AHL15. Previous analyses of genome-wide gene expression datasets using bioinformatics approaches have shown that neighboring genes in *Arabidopsis* are more co-expressed than random gene pairs (Williams and Bowles, 2004; Zhan *et al*., 2006; Chen *et al*., 2010; Wada *et al*., 2012; Yeaman, 2013; Kundu *et al*., 2017). Several mechanisms have been proposed to explain this phenomenon, such as gene duplications, shared promoters or common transcription factor binding motifs, but chromatin architecture has always been considered as the major source of co-expression of neighboring genes (Grob *et al*., 2014; Pombo and Dillon, 2015; Quintero-cadena and Sternberg, 2016). A recent genome-wide data analysis has excluded that co-regulation of neighboring genes in *Arabidopsis* is caused by gene duplications or the presence of common promoter motifs in neighboring genes (Kundu *et al*., 2017). Instead, co-regulation could be clearly correlated with local rearrangement of chromatin configuration (Kundu *et al*., 2017). Therefore, we suggest that the co-activation or co-repression of neighboring genes across genome by AHL15 is most likely also caused by extensive modulation of the chromatin configuration.

In animals, the contribution of higher-order chromatin organization to ageing processes has been well documented (Moraes, 2014; Chandra and Kirschner, 2016; Gorbunova and Seluanov, 2016). In contrast, the role of global organization of chromatin architecture in plant Ageing is not described yet. *Arabidopsis* adult leaves display a visible increase of chromatin compaction compared to juvenile leaves (Exner and Hennig, 2008), but the actual involvement of this chromatin compaction in the juvenile-to-adult transition has not been reported yet. Our results suggest that the global changes in gene expression after 4 hours of DEX-induced AHL15-GR activation coincide with rapid de-condensation of heterochromatin, thereby clearly linking plant ageing or tissue rejuvenation to respectively an increase or a reduction in the heterochromatin. Our results suggest that the extensive reprogramming of the transcriptome and the observed establishment of a long-term molecular memory following AHL15-GR activation might at least in part be caused by an extensive reorganization of the chromatin configuration.

We found that AHL15 represses several genes encoding components of the photosynthesis machinery. Recent studies have shown that the gradual increase in the sugar levels as a result of enhanced photosynthetic efficiency or an increase in leaf numbers promotes the juvenile-to-adult transition in *Arabidopsis*, tobacco, and tomato (Matsoukas *et al*., 2013; Yang *et al*., 2013; Yu *et al*., 2013). In contrast, a mutation in the *Arabidopsis CAO* gene that causes a low photosynthetic efficiency, was found to prolong the juvenile vegetative phase (Espine et al., 1999; Yu et al., 2013).

Unfortunately, how AHL15 alters the chromatin structure remains unclear. Detailed studies on the chromatin configuration by new approaches such as chromosome conformation capture technologies (Dekker et al., 2013), and comparative analysis of the putative binding sites of AHL15 with our transcriptome data are directions of future research that might provide more insight into the mode of action of this plant-specific AT-motif protein.

## Methods

### Plant material and RNA isolation and sequencing

The *Arabidopsis thaliana* (Col-0) plant line harboring the *35S::AHL15-GR* construct is described (Karami *et al*., 2020). The reporter lines and *H2B::H2B-GFP* (Fang and Spector, 2005) have been described previously. Seeds were surface sterilized (30 sec 70% ethanol, 10 minutes 1% chlorine, followed by washes in sterile water) and germinated after three days incubation at 4°C on MA medium (Masson and Paszkowski, 1992) containing 1% or no sucrose, and 0.7% agar at 21 °C and a 16 hours photoperiod. Following germination, 14 days old seedlings were transferred to soil and grown at 21 °C, 65% relative humidity, and long-day (LD: 16 hours photoperiod) condition.

For the transcriptome analysis (specifically), MA plates with 5 day old seedlings were flooded with water containing 1 ml ethanol per liter (mock), or with water to which dexamethasone (DEX, Sigma-Aldrich) dissolved in 1 ml ethanol was added to a final concentration of 20 μM. After 4 and 8 hours treatment, the shoot part of seedlings, including the shoot apex and cotyledons, was carefully separated from the hypocotyl and immediately frozen in liquid nitrogen and stored at −80 °C for RNA isolation.

Total RNA was extracted using a Qiagen RNeasy Plant Mini Kit, and the quality of the isolated RNA was validated using a nanodrop spectrophotometer (NanoDrop, ND-1000, Life Science). The isolated RNAs were reverse transcribed and sequenced on an Illumina HiSeq 2000 (100 nucleotides single reads). Three biological replicates were performed.

To test the effect of GA on AHL15-GR activation, 35 day old flowering *35S::AHL15-GR* plants were first sprayed with 20 μM DEX, followed 3 days later by spraying with 15 μM GA4 (Sigma-Aldrich).

### Quantification of expression levels and differential expression analysis

The quality control of all sequencing samples was carried out using FASTQC (version 0.10.1: http://www.bioinformatics.babraham.ac.uk/projects/fastqc/. Reference sequences and annotations for the *Arabidopsis* genome (TAIR10) were obtained from www.arabidopsis.org. To obtain expression levels the reads were aligned to the *Arabidopsis* genome sequence using Tophat2 (version 2.0.10) (Kim *et al*., 2013), using Bowtie2 (version 2.1.0) (Langmead and Salzberg, 2012) as the short read aligner at ‘very sensitive’ settings. Secondary alignments, i.e. alignments that meet Tophat’s criteria but are less likely to be correct than simultaneously reported primary alignments, were removed from the BAM files using SAMtools (version 0.1.18) (Li *et al*., 2009). Fragment alignment counts per transcript were determined from SAM alignment files using the Python package HTSeq-count (version 0.5.3p9) (Anders *et al*., 2014) with ‘strict’ settings to exclude reads aligning ambiguously with respect to annotated gene structures. Counts were summarized at the level of annotated genes, resulting in between 12.992.327 and 29.972.677 aligned fragments per sample. Read counts per annotated gene were normalized across all samples using the DESeq-like robust scaling factor (Anders and Huber, 2010) on reads per kilobase per million mapped reads (RPKM) values. For 13496 genes which had an expression value ≥ 10 in at least two samples, differential expression statistics were calculated using the R package edgeR (version 3.2.4) (Robinson *et al*., 2010).

### Gene ontology term analysis

Gene ontology (GO) term analysis for identification of enriched functional categories was performed using the agriGO single enrichment analysis tool (Du *et al*., 2010) (http://bioinformatics.cau.edu.cn/easygo/) with TAIR10 GO annotations. The MapMan software (Thimm *et al*., 2004) (http://mapman.gabipd.org/) was used to visualize pathways containing multiple genes with significant changes in expression.

### Quantitative real-time PCR analysis

RNA isolation was performed using a RevertAidTM Kit (Thermo Scientific). For qRT-PCR (qPCR), 1 μg of total RNA was used for cDNA synthesis with the iScript™ cDNA Synthesis Kit (BioRad). PCR was performed using the SYBR-Green PCR Master mix and amplification was run on a CFX96 thermal cycler (BioRad). The Pfaffl method was used to determine relative expression levels (Pfaffl, 2001). Expression was normalized using the *β-TUBULIN-6* gene. Three biological replicates were performed, with three technical replicates each. The primers used are described in Table 5.

**Table 5.**
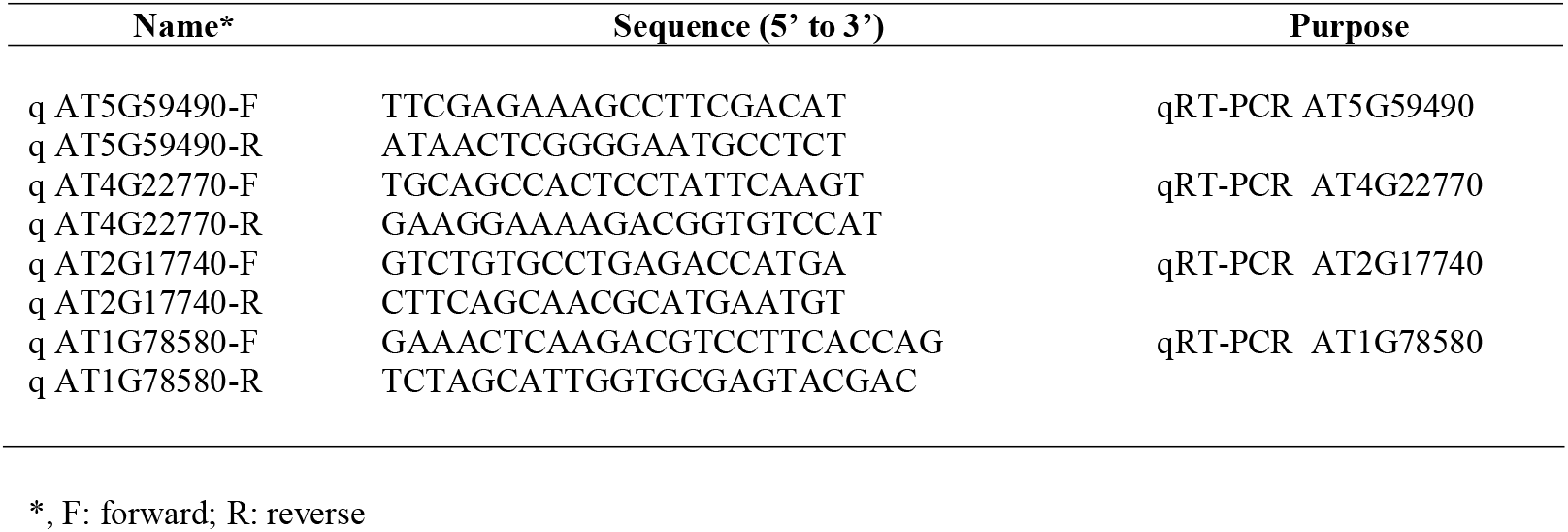
Sequences of DNA primers used for qRT-PCR (from 5’ to 3’)

### Microscopy

DAPI staining of nuclei was performed as described previously (Zanten *et al*., 2011). Samples were fixed in icecold Carnoys fixative (1:3 acetic acid:ethanol), then treated with enzymatic cell wall degrading solution containing 0.5% cellulose Onozuka R10 (Duchefa)), 0.25% macerozyme R10 (Duchefa), and 0.1% Triton X-100 for 1 h 30 min at room temperature. The samples were mounted with Vectashield (Vector laboratories) supplemented with 2 μg/ml 4’,6-diamidino-2-phenylindole (DAPI) before microscopic observation.

The heterochromatin phenotypes of the DAPI-stained leaf cells were recorded using a confocal laser scanning microscope (ZEISS-003-18533) using a 405 laser, a 350 nm LP excitation filter and a 425-475 nm BP emission filter. The H2B-GFP fusion protein was visualized using the same laser scanning microscope with a 534 laser, a 488 nm LP excitation filter and a 500-525 nm BP emission filters.

## Notes

### Competing Interest Statement

The authors have declared no competing interest.

